# Transgenic *Bax* gene efficiently induces lethality in mouse early embryos

**DOI:** 10.1101/2025.05.26.656236

**Authors:** Yuzuki Goto, Takuto Yamamoto, Masahiro Sakata, Satoshi Mashiko, Daiki Shikata, Shinnosuke Honda, Naojiro Minami, Shuntaro Ikeda

**Affiliations:** Laboratory of Reproductive Biology, Graduate School of Agriculture, Kyoto University, Kyoto 606-8502, Japan

**Keywords:** *Bax*, mouse early embryos, apoptosis, Tet-On system

## Abstract

Apoptosis is an essential physiological process involved in embryonic development, immune responses, and tissue homeostasis. Despite many studies on pro-apoptotic genes, few reports have directly compared the lethality-inducing potential between them under identical conditions. In this study, we evaluated the lethality-inducing potential of three representative pro-apoptotic genes, *Bax*, *Casp3*, and *Casp9*, in mouse early embryos under defined conditions using the doxycycline (Dox)-inducible tetracycline-regulated gene expression system (Tet-On system) in combination with the PiggyBac transposon system. All genes were transcriptionally induced by Dox, and *Bax* showed the strongest lethal effect, followed by *Casp9,* while *Casp3* did not show any effect. Notably, *Bax* expression severely impaired blastocyst formation and led to the intense accumulation of the DNA damage marker γH2AX. These findings suggest that introducing upstream apoptotic regulators leads to the more efficient and widespread activation of the apoptotic cascade. Additionally, an unexpected Dox-dependent increase in the expression of reverse tetracycline-controlled transactivator, which is typically driven by a constitutive promoter, was observed, raising the possibility of unanticipated regulatory mechanisms within the Tet-On system. Overall, this study is expected to contribute to a deeper understanding of apoptotic mechanisms and future advancements in regenerative medicine, reproductive engineering, and cancer research.

## Introduction

Apoptosis is a fundamental and essential physiological process in multicellular organisms, playing critical roles in embryonic development, immune responses, and the maintenance of tissue homeostasis. It is a highly regulated mechanism that eliminates unnecessary, aged, or damaged cells without inducing an inflammatory response [1–4].

Apoptosis is triggered by two major signaling pathways, the extrinsic and intrinsic pathways, both of which ultimately lead to the activation of executioner caspases, such as caspase-3 and caspase-7, resulting in cell death [1–6]. The extrinsic pathway is initiated when extracellular death signals, such as Fas ligand or TRAIL, bind to death receptors on the cell membrane, leading to the activation of caspase-8 via FADD and subsequently resulting in the activation of executioner caspases [7, 8]. In contrast, the intrinsic pathway is activated in response to intracellular stress signals, such as DNA damage, oxidative stress, or loss of growth factor [9]. In this pathway, B-cell lymphoma-2 (Bcl-2) family members such as Bax and Bak induce mitochondrial outer membrane permeabilization (MOMP), leading to the release of cytochrome c into the cytosol [10]. Released cytochrome c binds to apoptotic protease activating factor 1 (Apaf-1), which subsequently activates caspase-9. Activated caspase-9 then triggers the activation of executioner caspases [9].

These recent advances in our understanding of the molecular mechanisms of apoptosis have led to increased efforts to intentionally “utilize” this system. Bax, caspase-3, and caspase-9 are central effector molecules that play crucial roles in apoptotic pathways; therefore, they have been frequently investigated in basic and applied studies [11–20]. Specifically, the overexpression of these genes to induce selective apoptosis in target cells has been widely explored. For example, the overexpression of Bax or caspase-3 has been shown to effectively induce tumor cell death, positioning this approach as a promising strategy for cancer therapy [21, 22]. In addition, caspase-9 overexpression has been demonstrated to induce apoptosis in primary cultures of anterior pituitary cells and HeLa tumor cells, suggesting its potential utility as a model toward understanding the pathological mechanisms of neurodegenerative or malignant diseases and toward developing new therapeutic strategies [23].

Thus, the strategic use of apoptotic pathways holds promising applications in the medical fields including cancer therapy and neurodegenerative diseases. Consequently, the ability to effectively induce apoptosis has become a key focus. In this context, identifying which pro-apoptotic genes can most efficiently trigger cell death is important for expanding its medical and related applications as well as for advancing our fundamental understanding of apoptotic mechanisms. However, few studies have directly compared the apoptotic potency of these genes under identical experimental conditions.

Therefore, we evaluated the lethality-inducing potential of three representative pro-apoptotic genes: *Bax*, *Casp3*, and *Casp9*. To achieve this, we employed the doxycycline (Dox)-inducible tetracycline-regulated gene expression system (Tet-On system), which enables precise control of gene expression in mammalian cells [24, 25], together with the PiggyBac transposon system, a highly efficient and easy to use method for gene integration in mammalian cells [26–29]. By introducing either *Bax*, *Casp3*, or *Casp9* into mouse zygotes using these two tools, and precisely controlling the expression of each pro-apoptotic gene in a Dox-dependent manner, we performed a comparative evaluation of their ability to induce cell lethality under identical conditions.

This study, using mammalian zygotes as a model, provides a novel framework for comparing the lethality-inducing potential of pro-apoptotic genes under identical conditions, thereby offering new insights into their apoptosis-inducing potential during the early developmental stages of mammalian embryos. In conventional models, such as cancer cells, the genetic background is often heterogeneous due to their differentiated state, and the genome is unstable, leading to cell cycle abnormalities. Additionally, the anti-apoptotic activity of cancer cells [30] limits our ability to compare and evaluate the effects of pro-apoptotic genes. In contrast, zygotes possess pluripotency and undergo synchronous and well-ordered developmental processes, making them a suitable model for accurately evaluating the physiological effects of the introduced genes. By deepening our fundamental understanding of pro-apoptotic gene function, this research not only contributes to basic biology but also lays the groundwork for future applications such as reproductive engineering. Ultimately, these findings may support the development of foundational technologies for the medical fields including cancer therapy.

## Materials and Methods

### In vitro fertilization (IVF)

Female ICR mice (8–12 weeks old; Japan SLC, Shizuoka, Japan) were superovulated by intraperitoneal injection of 7.5 IU equine chorionic gonadotropin (eCG; ASKA Animal Health, Tokyo, Japan), followed by 7.5 IU human chorionic gonadotropin (hCG; ASKA Animal Health) 46–48 h later. At 14–16 h after hCG injection, the mice were euthanized by cervical dislocation, and cumulus oocyte complexes (COCs) were collected and placed in human tubal fluid (HTF) medium supplemented with 4 mg/mL bovine serum albumin (BSA; Sigma-Aldrich, St. Louis, MO, USA) [31]. Male ICR mice (13–16 weeks old; Japan SLC) were euthanized by cervical dislocation, and spermatozoa were collected and cultured in HTF medium for at least 1 h for capacitation. The sperm suspension was then added to the fertilization droplets containing COCs at a final concentration of 1.0 × 10^6^ cells/mL, and the gametes were co-incubated for 2–3 h at 37°C in a humidified atmosphere containing 5% CO_2_. After fertilization, the embryos were washed three times in KSOM supplemented with amino acids [32] and 1 mg/mL BSA to remove cumulus cells and excess sperm. Morphologically normal zygotes were used for subsequent microinjection procedures.

### Plasmid construction and in vitro transcription (IVT)of hyPBase mRNA

The coding sequence (CDS) of *Bax* was amplified from total cDNA extracted from mouse ovaries by nested PCR using KOD-Plus polymerase (Toyobo, Osaka, Japan). Outer primers targeting the flanking regions and nested primers specific to the target region were designed to perform a two-step amplification. The amplified product was digested with *Not*I (New England Biolabs, Ipswich, MA, USA) and *Xba*I (New England Biolabs) and cloned into a pBluescript II SK (−) plasmid (Agilent Technologies, Santa Clara, CA, USA), which had been inserted with a FLAG tag (DYKDDDDK) and a linker sequence downstream of the T7 promoter (pBS-*Bax*). Ligation was performed using a Ligation High Kit (Toyobo). From this point onward, DNA fragments were assembled to generate each construct by using Gibson Assembly Master Mix (New England Biolabs). The pTet-One vector from the Tet-One Inducible Expression System (Takara Bio, Kusatsu, Japan) and *Bax* from pBS-*Bax* were amplified by PCR using KOD-Plus Neo polymerase (Toyobo), and assembled into pTet-One-*Bax*. A pAcGFP-membrane (mem) vector (Clontech, Mountain View, CA, USA) and an IRES fragment were used to construct IRES-mem-AcGFP. The pTet-One-*Bax* and IRES-mem-AcGFP fragments were amplified with KOD-Plus Neo, and IRES-mem-AcGFP was fused downstream of the Tet-On 3G (reverse tetracycline-controlled transactivator, rtTA) gene to generate pTet-One-*Bax*-IRES-mem-AcGFP. The inverted terminal repeat-flanked backbone, excluding the CAG-TagRFP sequence from pPB-CAG-TagRFP [33], and the genetic cassette of pTet-One-*Bax*-IRES-mem-AcGFP were each PCR amplified with KOD-Plus Neo and assembled to generate pPB-Tet-One-*Bax*-IRES-mem-AcGFP. The CDS of *Casp3* and *Casp9* were PCR amplified from total cDNA derived from mouse brain using KOD-Plus Neo and cloned into pBluescript II SK (−). Subsequently, the pPB-Tet-One-*Bax*-IRES-mem-AcGFP sequence, excluding the *Bax* region, and *Casp3* or *Casp9* sequence were each PCR amplified with KOD-Plus Neo and fused to construct pPB-Tet-One-*Casp3*-IRES-mem-AcGFP and pPB-Tet-One-*Casp9*-IRES-mem-AcGFP, respectively. To generate pPB-Tet-One-(no gene)-IRES-mem-AcGFP, the pPB-Tet-One-*Casp3*-IRES-mem-AcGFP sequence, excluding the *Casp3* region, was amplified using a KOD-Plus-Mutagenesis Kit (Toyobo) and self-ligated. Capped and polyadenylated hyPBase mRNA was synthesized *in vitro* using an mMESSAGE mMACHINE T7 Ultra Kit (Thermo Fisher Scientific, Waltham, MA, USA) from pCAG-hyPBase [29], and purified with an RNeasy Mini Kit (Qiagen, Hilden, Germany). The resulting mRNA was resuspended in nuclease-free water. The primer sequences used for plasmid construction and IVT are listed in Supplementary Table 1.

### Plasmid microinjection, Dox treatment, and embryo culture

Approximately 3–5 pL of a mixture containing 30 ng/µL hyPBase mRNA and 30 ng/µL of either pPB-Tet-One-(no gene)-IRES-mem-AcGFP, pPB-Tet-One-*Bax*-IRES-mem-AcGFP, pPB-Tet-One-*Casp3*-IRES-mem-AcGFP, or pPB-Tet-One-*Casp9*-IRES-mem-AcGFP was microinjected into the cytoplasm of each zygote at 3–6 hours post-insemination (hpi) [27]. After microinjection, morphologically normal embryos with two pronuclei were collected from each group and divided for culture in KSOM supplemented with (Dox+) or without (Dox−) 100 ng/mL Dox [34] from 6 to 96 hpi. To assess the effect of Dox on embryonic development, uninjected embryos were also cultured under the same conditions. All embryos were incubated at 37°C in a humidified atmosphere containing 5% CO_2_.

### RNA extraction and reverse transcription-quantitative PCR (RT-qPCR)

By using a QuantAccuracy™ RT-RamDA™ cDNA Synthesis Kit (Toyobo), total RNA was extracted from 5 embryos in each experimental group and reverse transcribed into cDNA. Quantitative PCR was carried out using the synthesized cDNA as a template with gene-specific primers and KOD SYBR qPCR Mix (Toyobo). Amplification and quantification of gene transcripts were conducted following a previously established protocol [35]. *H2afz* was used as an internal control. Relative expression levels were determined using the comparative Ct method (2^−ΔΔCt^) [36]. The primer sequences used for the RT-qPCR are listed in Supplementary Table1.

### Immunofluorescence and analysis

To evaluate apoptosis, the embryos were fixed with 4% paraformaldehyde in phosphate-buffered saline (PBS) for 20 min at 28°C after removal of the zona pellucida with acid Tyrode’s solution (pH 2.5). Subsequently, the embryos were permeabilized in 0.5% Triton X-100 (Sigma-Aldrich) in PBS for 40 min at 28°C. After permeabilization, the embryos were blocked in PBS containing 1.5% BSA, 0.2% sodium azide, and 0.02% Tween 20 (blocking buffer) for 1 h at 28°C, and then incubated overnight at 4°C with a mouse anti-H2A.X (Ser139) antibody (1:200 dilution; 613401; BioLegend, San Diego, CA, USA). The embryos were washed in blocking buffer and incubated for 1 h at 28°C with an Alexa Fluor 647 donkey anti-mouse IgG (H+L) secondary antibody (1:500 dilution; Thermo Fisher Scientific). After washing, the embryos were stained with 10 μg/mL Hoechst 33342 (Sigma-Aldrich) in blocking buffer for 20 min at 28°C. Finally, the embryos were mounted on glass slides and observed using a fluorescence microscope (IX73; Olympus, Tokyo, Japan). The fluorescence intensities of γH2AX in embryos was quantified using ImageJ software (National Institutes of Health, Bethesda, MD). First, the embryo region in the image was selected, and the mean fluorescence intensity (Mean Gray Value) within this region was measured. Simultaneously, the area (Area) of the selected region was measured, and the integrated density (Integrated Density) was calculated by multiplying the mean intensity by the area.

### Statistical analysis

Developmental rates were analyzed using a chi-square test. Differences in GFP and γH2AX fluorescence intensities were analyzed using Student’s *t*-test (or the Mann–Whitney *U* test if the data were not normally distributed). The experimental groups were as follows: (1) non-injected (Non-injected) embryos; (2) gene-lacking construct-injected (Control) embryos; (3) *Bax*-injected (*Bax*) embryos; (4) *Casp3*-injected (*Casp3*) embryos; and (5) *Casp9*-injected (*Casp9*) embryos. For each group, statistical comparisons were made between the Dox− and Dox+ treatments, and *P*-values < 0.05 were considered statistically significant. This approach was chosen to evaluate clearly the impact of Dox-induced gene expression on development, while distinguishing it from the effects of Dox treatment itself and the effects of injection.

### Ethical approval for the use of animals

All animal experiments were conducted with approval from the Animal Research Committee of Kyoto University (approval nos.: R3–17, R4–17, R5–17, and R6–17) and in strict compliance with the committee’s ethical guidelines.

## Results

### Overexpression of pro-apoptotic genes in preimplantation embryos

To induce the overexpression of pro-apoptotic genes, plasmids carrying each gene (*Bax*, *Casp3*, or *Casp9*) were co-injected with hyPBase mRNA into the cytoplasm of mouse zygotes at 3–6 hpi, and the embryos were cultured in KSOM supplemented with (Dox+) or without (Dox−) 100 ng/mL Dox until 96 hpi (Fig. 1B). The expression of each pro-apoptotic transgene was analyzed at the blastocyst stage. The expression of all transgenes was increased in embryos in Dox+ compared with those in Dox− (Fig. 2A). These results confirm that the Tet-On system introduced via the PiggyBac transposon system functioned properly, enabling the Dox-inducible overexpression of pro-apoptotic transgenes.

**Fig. 1.**
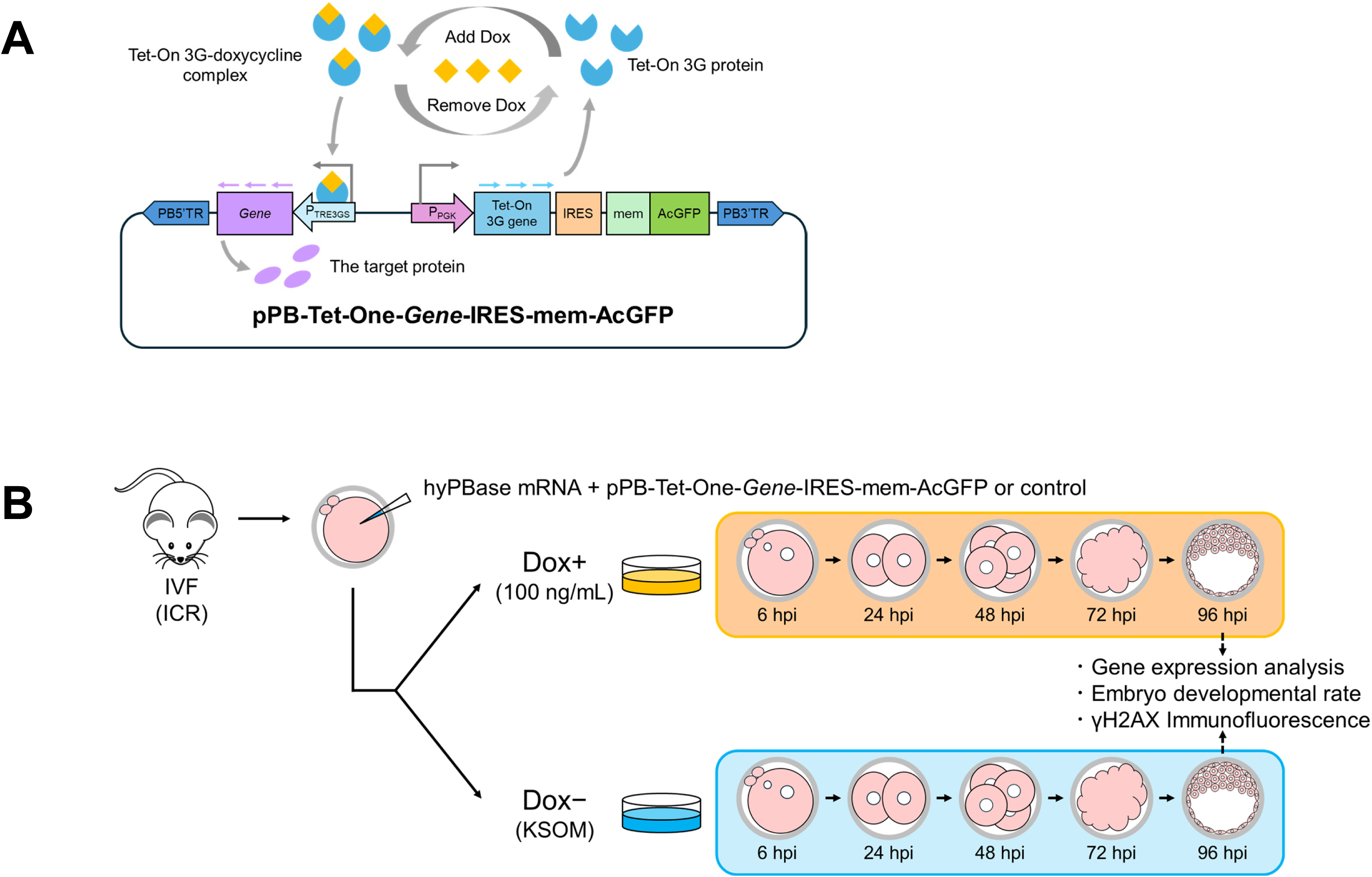
Overview of the experimental procedure. (A) Schematic diagram of a PiggyBac-based all-in-one vector system designed for Dox-inducible gene expression. The construct contains a gene of interest under the control of the TRE3GS promoter, which is activated upon Dox-inducible binding of the Tet-On 3G (rtTA) protein. The Tet-On 3G gene is constitutively expressed under the PGK promoter. The IRES-mem-AcGFP cassette allows for bicistronic expression. PB5′TR and PB3′TR are the terminal repeats required for PiggyBac transposase. (B) Schematic diagram of the experimental design. The embryos were microinjected with hyPBase mRNA and each plasmid, cultured in the presence or absence of Dox, and analyzed for gene expression, developmental rate, and γH2AX signal.

**Fig. 2.**
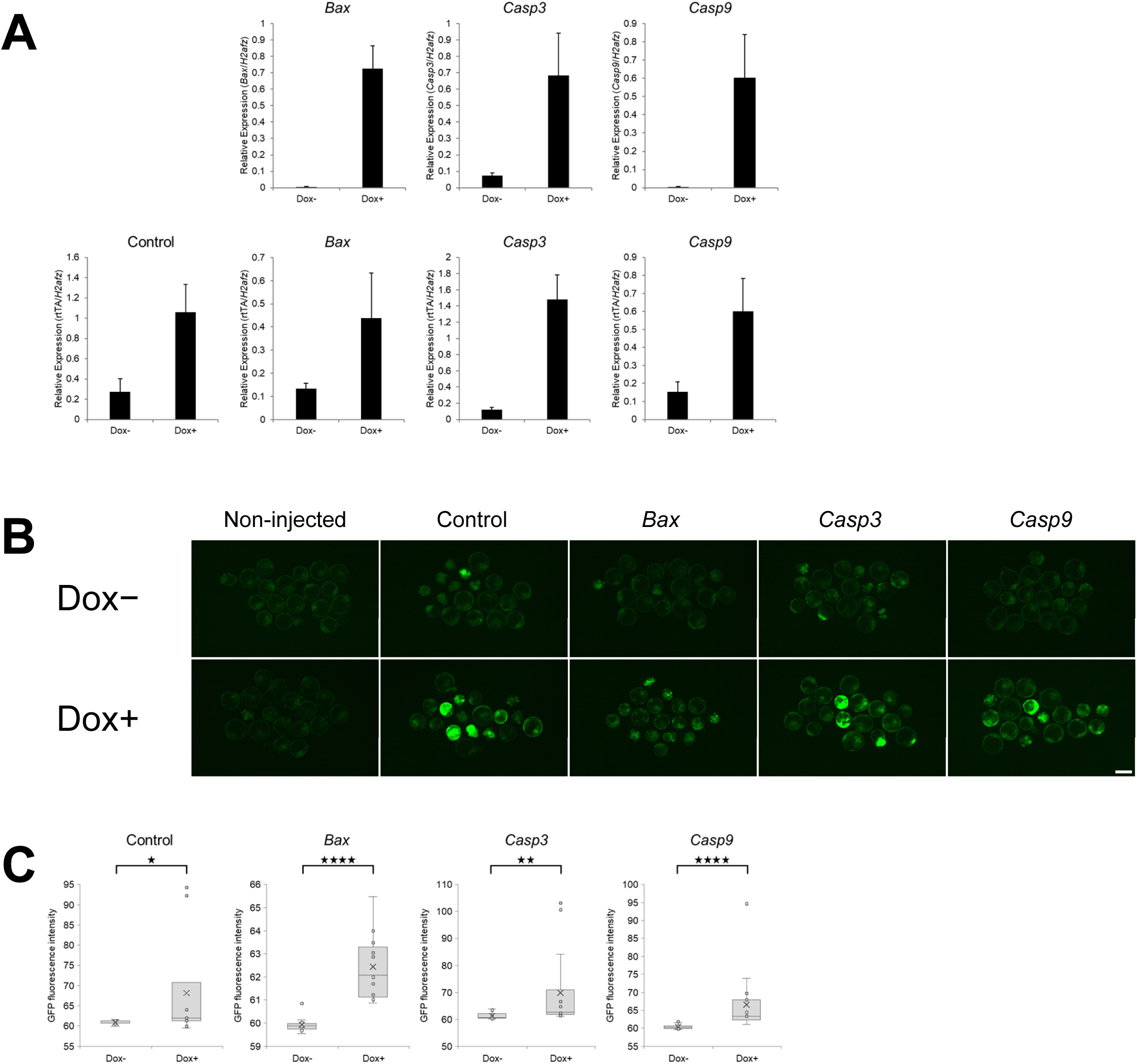
Verification of gene introduction and expression levels. The experimental groups were as follows: (1) non-injected (Non-injected) embryos; (2) gene-lacking construct-injected (Control) embryos; (3) *Bax*-injected (*Bax*) embryos; (4) *Casp3*-injected (*Casp3*) embryos; and (5) *Casp9*-injected (*Casp9*) embryos. (A) RT-qPCR of each pro-apoptotic gene (*Bax*, *Casp3*, and *Casp9*) and rtTA in each group was performed using 5 embryos at the blastocyst (96 hpi) stage. Gene expression levels were normalized to *H2afz* as an internal control. Data are expressed as the mean ± standard error of the mean (*n* = 3). (B) GFP expression in mouse preimplantation embryos in each experimental group at 96 hpi. Scale bar, 100 µm. (C) GFP fluorescence intensity. Control (Dox−), *n* = 11; Control (Dox+), *n* = 11; *Bax* (Dox−), *n* = 13; *Bax* (Dox+), *n* = 13; *Casp3* (Dox−), *n* = 14; *Casp3* (Dox+), *n* = 14; *Casp9* (Dox−), *n* = 15; *Casp9* (Dox+), *n* = 15. ^★^*P* < 0.05, ^★★^*P* < 0.01, ^★★★★^*P* < 0.0001, Mann–Whitney *U* test.

Simultaneously, we attempted to verify the presence of the knock-in by monitoring GFP fluorescence, which was placed downstream of the Tet-On 3G (rtTA) gene regardless of Dox treatment (Fig. 1A). Because rtTA gene expression is considered to be constitutive regardless of Dox treatment [24, 25], comparable levels of GFP fluorescence were expected between the Dox−and Dox+ groups. However, surprisingly, GFP fluorescence intensity was significantly higher in Dox+ embryos than in Dox− embryos in all groups: Control, *Bax*, *Casp3*, and *Casp9* (Fig. 2B, C). To confirm this unexpected observation, we quantified rtTA gene expression in the above groups. Consequently, rtTA gene expression levels were consistently elevated in Dox+ embryos in all groups (Fig. 2A). This result deviates from previously established findings [24, 25].

### Bax overexpression effectively induces apoptosis in mouse embryos

The effects of apoptosis-inducing genes on embryonic development were monitored in each group up to 96 hpi. Among the tested genes, the Dox-induced overexpression of *Bax* led to a significant reduction in the proportion of embryos reaching the blastocyst stage, indicating the strongest inhibitory effect on development among these three genes. The Dox-induced overexpression of *Casp9* inhibited embryonic development to a certain degree, although its inhibitory effect was less pronounced than that of *Bax*. In contrast, the Dox-induced overexpression of *Casp3* had no effect on the developmental rate, showing similar outcomes to the Control embryos with no apparent developmental inhibition (Fig. 3, Table 1). Furthermore, in the Non-injected embryos, the developmental rate was similar between the Dox− and Dox+ conditions, indicating that Dox at a concentration of 100 ng/mL does not adversely affect embryonic development.

**Fig. 3.**
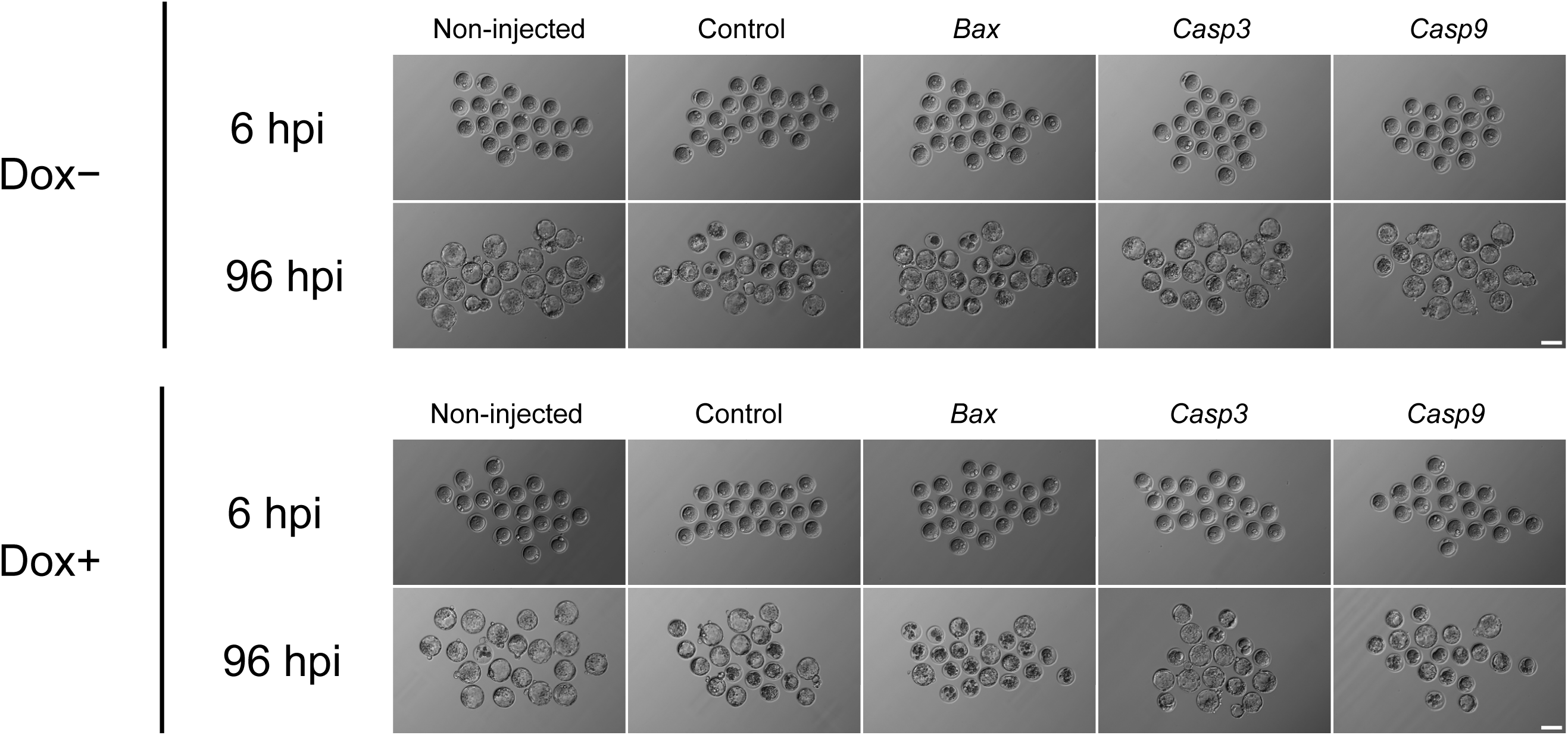
Embryonic development and apoptosis induction at the blastocyst stage. Developmental morphology of the embryos at the start (6 hpi) and end (96 hpi) of culture in Dox− or Dox+. Scale bar, 100 μm.

**Table 1.** Developmental rate in the absence (−) or presence (+) of Dox for each treatment group ns, not significant, ^★★^*P* < 0.01, ^★★★★^*P* < 0.0001, chi-square test in each treatment.

| Treatment | Dox<br>(–/+) | No. of embryos with two<br>pronuclei | No. of<br>blastocysts | Ratio of blastocysts per<br>fertilized embryos (%) |  |
| --- | --- | --- | --- | --- | --- |
| Non-injected | Dox– | 50 | 49 | 98.0 | ] n.s. |
| Non-injected | Dox+ | 88 | 83 | 94.3 |  |
| Control | Dox– | 63 | 49 | 77.8 | ] n.s. |
| Control | Dox+ | 94 | 68 | 72.3 |  |
| *Bax* | Dox– | 72 | 60 | 83.3 | ] **** |
| *Bax* | Dox+ | 88 | 45 | 51.1 |  |
| *Casp3* | Dox– | 69 | 52 | 75.4 | ] n.s. |
| *Casp3* | Dox+ | 76 | 60 | 78.9 |  |
| *Casp9* | Dox– | 53 | 45 | 84.9 | ] ** |
| *Casp9* | Dox+ | 96 | 62 | 64.6 |  |

To evaluate the cellular responses induced by the expression of these pro-apoptotic genes, we assessed DNA damage in blastocyst stage embryos using γH2AX immunostaining as a marker for apoptosis. Dox-induced *Bax* overexpression resulted in a robust increase in the γH2AX signal, reflecting widespread DNA damage. A moderate increase was observed in the *Casp9* group. In contrast, the *Casp3* group did not exhibit any significant difference in γH2AX signal intensity between the Dox− and Dox+ conditions, similar to the Non-injected and Control embryos (Fig. 4A, B).

**Fig. 4.**
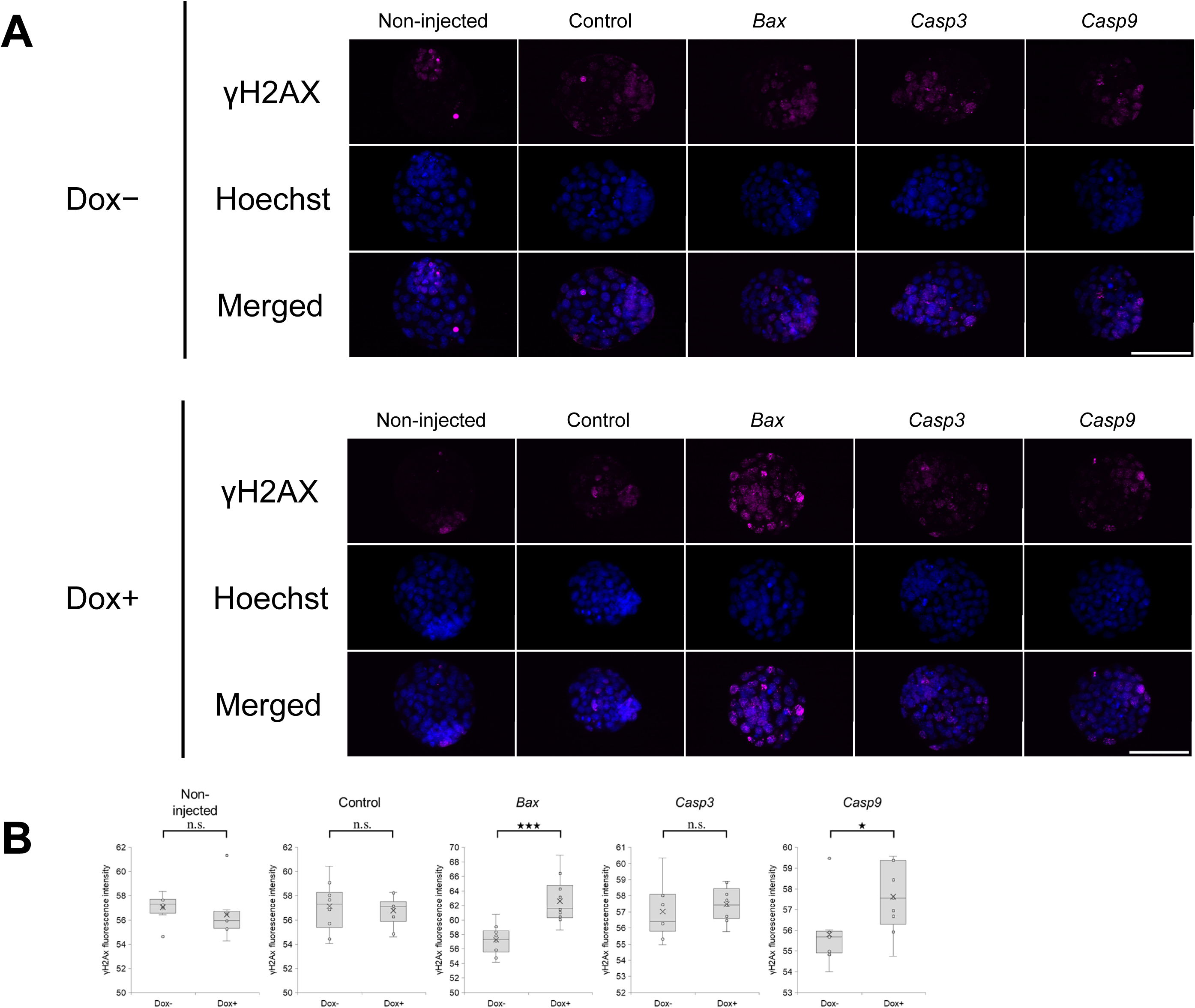
Detection of apoptosis by immunofluorescence. (A) DNA damage in mouse preimplantation embryos at the blastocyst stage (96 hpi) detected by γH2AX immunostaining. The nuclei were counterstained using Hoechst 33342. Scale bar, 100 μm. (B) γH2AX immunofluorescence intensity. Non-injected (Dox−), *n* = 8; Non-injected (Dox+), *n* = 8; Control (Dox−), *n* = 10; Control (Dox+), *n* = 10; *Bax* (Dox−), *n* = 10; *Bax* (Dox+), *n* = 10; *Casp3* (Dox−), *n* = 9; *Casp3* (Dox+), *n* = 9; *Casp9* (Dox−), *n* = 9; *Casp9* (Dox+), *n* = 9. ns, not significant, ^★^*P* < 0.05, ^★★★^*P* < 0.001, unpaired *t*-test (or the Mann–Whitney *U* test for non-normally distributed data).

## Discussion

This study was conducted to compare the lethality-inducing potential of three representative pro-apoptotic genes—*Bax*, *Casp3*, and *Casp9*—in mouse early embryos under identical conditions, by combining the Tet-On system with transgenesis using the PiggyBac transposon system. As a result, all three genes exhibited Dox-dependent upregulation at the transcriptional level; however, their effects on embryonic development and apoptosis varied markedly.

Among the three genes tested, the induction of *Bax* expression most strongly inhibited embryonic development and markedly increased γH2AX signals, suggesting robust DNA damage and the induction of apoptosis. This may reflect the fact that Bax is an upstream molecule that directly initiates apoptosis by inducing MOMP [10], a key triggering event in the apoptotic pathway, consistent with a previous finding that Bax alone can trigger apoptosis in mammalian cells [37].

In contrast, caspase-3 and caspase-9 function downstream of Bax and require upstream activators for their activation. This may be a reason for their weaker inhibitory effects on embryonic development upon induction. In the case of *Casp3*, no impact was shown. The induction of *Casp9* expression resulted in a certain degree of developmental inhibition, possibly because it acts upstream of caspase-3 and can activate not only caspase-3 but also caspase-7 and, indirectly, caspase-6 [38, 39], resulting in a stronger effect than that of *Casp3* alone.

Interestingly, in all of the experimental groups in which the Tet-On system was introduced, the addition of Dox resulted in increased rtTA mRNA levels and GFP fluorescence. This was an unexpected result, given that rtTA is typically driven by a constitutive promoter and its expression is Dox-independent [24, 25]. The improved Tet-On system used in this study is an all-in-one vector [40, 41]. Therefore, feedback or crosstalk between its components may be involved, although the details remain unclear. Further analysis of the molecular mechanism is needed.

A limitation of this study is that gene expression was assessed only at the mRNA level, without direct evaluation of protein translation or the activation states of caspases. Furthermore, while γH2AX was used as a marker to evaluate apoptosis, incorporating additional assays, such as the TUNEL assay or immunofluorescence of active caspases, would provide more robust and complementary evidence.

In summary, this study demonstrated that *Bax* exhibits the strongest lethality-inducing potential in mouse early embryos, followed by *Casp9*, while *Casp3* did not show any lethal effect. These results suggest that the further upstream an introduced molecule is, the wider range of the apoptotic cascade it can efficiently activate. In contrast, *Casp9* and *Casp3* are highly dependent on the activation conditions, indicating the need for optimization for their effective use. Additionally, an unexpected behavior was observed in the Tet-On system, highlighting the need for further investigation. Overall, these findings enhance our understanding of apoptotic mechanisms and provide a foundation for future applications in regenerative medicine, reproductive engineering, and cancer research.

## Figure legends

**Supplementary Table 1.**
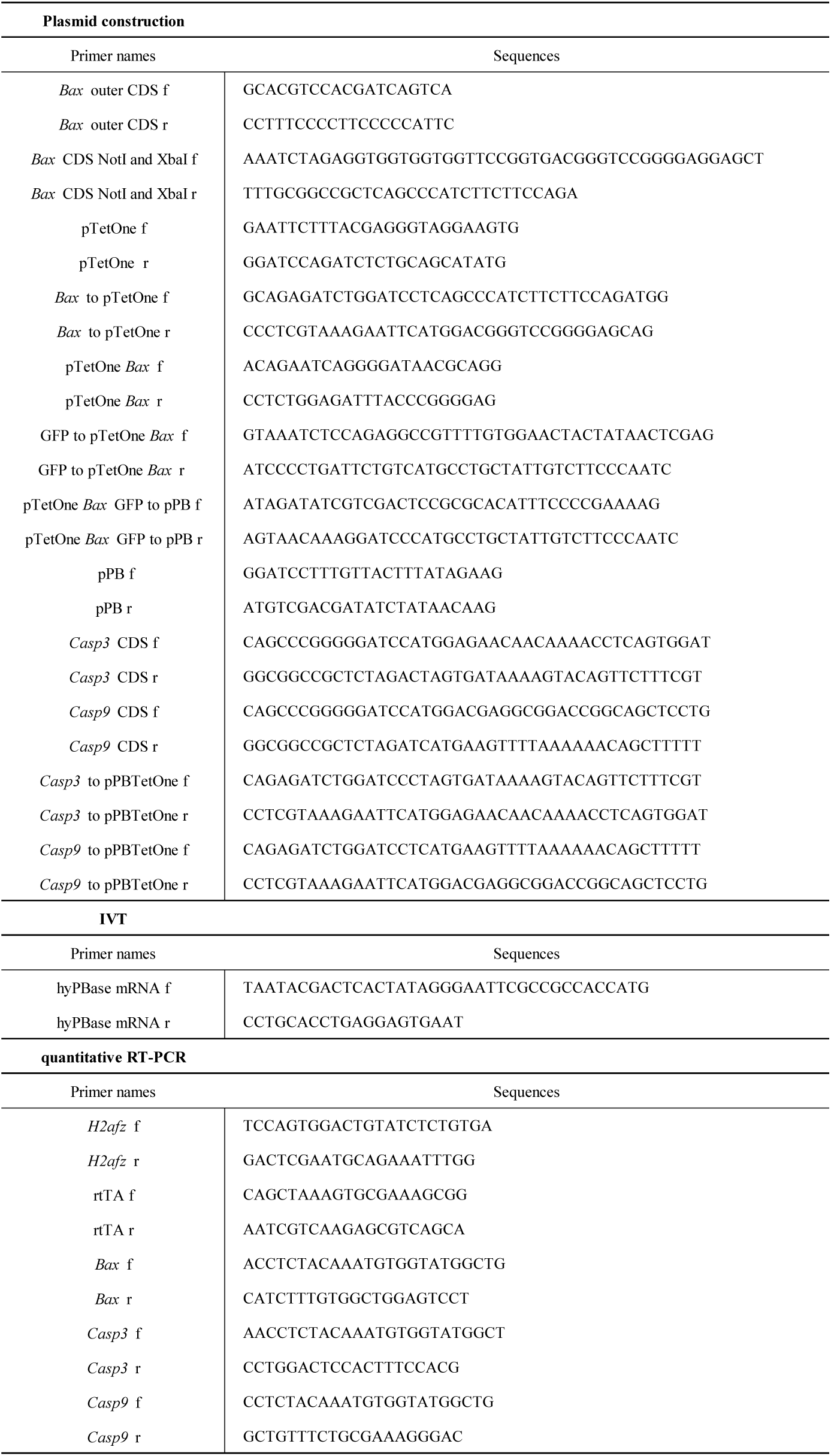
Primer sequences.

## References

1. Danial, N. N., Korsmeyer, S. J. Cell Death: Critical Control Points. Cell 2004; 116: 205–219.

2. D’Arcy, M. S. Cell death: a review of the major forms of apoptosis, necrosis and autophagy. Cell Biology International 2019; 43: 582–592.

3. Elmore, S. Apoptosis: A Review of Programmed Cell Death. Toxicologic Pathology 2007; 35: 495–516.

4. Nagata, S. Apoptosis and Clearance of Apoptotic Cells. Annual Review of Immunology 2018; 36: 489–517.

5. Green, D. R., Llambi, F. Cell Death Signaling. Cold Spring Harbor Perspectives in Biology 2015; 7: a006080.

6. Kroemer, G., Galluzzi, L., Brenner, C. Mitochondrial membrane permeabilization in cell death. Physiological Reviews 2007; 87: 99–163.

7. Ashkenazi, A., Dixit, V. M. Death receptors: Signaling and modulation. Science 1998; 281: 1305–1308.

8. Nair, P., Lu, M., Petersen, S., Ashkenazi, A. Apoptosis initiation through the cell-extrinsic pathway. Methods in Enzymology 2014; 544: 99–128.

9. Tait, S. W. G., Green, D. R. Mitochondria and cell death: outer membrane permeabilization and beyond. Nature Reviews Molecular Cell Biology 2010 11:9 2010; 11: 621–632.

10. Youle, R. J., Strasser, A. The BCL-2 protein family: opposing activities that mediate cell death. Nature Reviews Molecular Cell Biology 2008 9:1 2008; 9: 47–59.

11. Brady, H. J. M., Gil-Gómez, G. Molecules in focus Bax. The pro-apoptotic Bcl-2 family member, Bax. The International Journal of Biochemistry & Cell Biology 1998; 30: 647–650.

12. Brentnall, M., Rodriguez-Menocal, L., De Guevara, R. L., Cepero, E., Boise, L. H. Caspase-9, caspase-3 and caspase-7 have distinct roles during intrinsic apoptosis. BMC Cell Biology 2013; 14: 1–9.

13. Jänicke, R. U., Sprengart, M. L., Wati, M. R., Porter, A. G. Caspase-3 is required for DNA fragmentation and morphological changes associated with apoptosis. Journal of Biological Chemistry 1998; 273: 9357–9360.

14. Jiang, M., Qi, L., Li, L., Li, Y. The caspase-3/GSDME signal pathway as a switch between apoptosis and pyroptosis in cancer. Cell Death Discovery 2020; 6: 1–11.

15. Kuida, K. Caspase-9. The International Journal of Biochemistry & Cell Biology 2000; 32: 121–124.

16. Li, P., Zhou, L., Zhao, T., Liu, X., Zhang, P., Liu, Y., Zheng, X., Li, Q. Caspase-9: structure, mechanisms and clinical application. Oncotarget 2017; 8: 23996.

17. Renault, T. T., Manon, S. Bax: Addressed to kill. Biochimie 2011; 93: 1379–1391.

18. Würstle, M. L., Laussmann, M. A., Rehm, M. The central role of initiator caspase-9 in apoptosis signal transduction and the regulation of its activation and activity on the apoptosome. Experimental Cell Research 2012; 318: 1213–1220.

19. Yin, C., Knudson, C. M., Korsmeyer, S. J., Van Dyke, T. Bax suppresses tumorigenesis and stimulates apoptosis in vivo. Nature 1997; 385: 637–640.

20. Zhang, L., Yu, J., Park, B. H., Kinzler, K. W., Vogelstein, B. Role of BAX in the Apoptotic Response to Anticancer Agents. Science 2000; 290: 989–992.

21. Melis, M. H. M., Simpson, K. L., Dovedi, S. J., Welman, A., MacFarlane, M., Dive, C., Honeychurch, J., Illidge, T. M. Sustained tumour eradication after induced caspase-3 activation and synchronous tumour apoptosis requires an intact host immune response. Cell Death & Differentiation 2013 20:5 2013; 20: 765–773.

22. Zheng, J. Y., Yang, G. S., Wang, W. Z., Li, J., Li, K. Z., Guan, W. X., Wang, W. L. Overexpression of Bax induces apoptosis and enhances drug sensitivity of hepatocellular cancer-9204 cells. World Journal of Gastroenterology : WJG 2005; 11: 3498.

23. Druškovič, M., Šuput, D., Milisav, I. Overexpression of Caspase-9 Triggers Its Activation and Apoptosis in Vitro. Croatian medical journal 2006; 47: 832. Retrieved from https://pmc.ncbi.nlm.nih.gov/articles/PMC2080483/

24. Gossen, M., Bujard, H. Tight control of gene expression in mammalian cells by tetracycline-responsive promoters. Proceedings of the National Academy of Sciences of the United States of America 1992; 89: 5547–5551.

25. Goverdhana, S., Puntel, M., Xiong, W., Zirger, J. M., Barcia, C., Curtin, J. F., Soffer, E. B., Mondkar, S., King, G. D., Hu, J., Sciascia, S. A., Candolfi, M., Greengold, D. S., Lowenstein, P. R., Castro, M. G. Regulatable gene expression systems for gene therapy applications: progress and future challenges. Molecular Therapy 2005; 12: 189–211.

26. Ding, S., Wu, X., Li, G., Han, M., Zhuang, Y., Xu, T. Efficient transposition of the piggyBac (PB) transposon in mammalian cells and mice. Cell 2005; 122: 473–483.

27. Suzuki, S., Tsukiyama, T., Kaneko, T., Imai, H., Minami, N. A hyperactive piggyBac transposon system is an easy-to-implement method for introducing foreign genes into mouse preimplantation embryos. The Journal of reproduction and development 2015; 61: 241–244.

28. Wu, S. C. Y., Meir, Y. J. J., Coates, C. J., Handler, A. M., Pelczar, P., Moisyadi, S., Kaminski, J. M. piggyBac is a flexible and highly active transposon as compared to Sleeping Beauty, Tol2, and Mos1 in mammalian cells. Proceedings of the National Academy of Sciences of the United States of America 2006; 103: 15008–15013.

29. Yusa, K., Zhou, L., Li, M. A., Bradley, A., Craig, N. L. A hyperactive piggyBac transposase for mammalian applications. Proceedings of the National Academy of Sciences of the United States of America 2011; 108: 1531–1536.

30. Hanahan, D., Weinberg, R. A. Hallmarks of cancer: The next generation. Cell 2011; 144: 646–674.

31. Minami, N., Sasaki, K., Aizawa, A., Miyamoto, M., Imai, H. Analysis of gene expression in mouse 2-cell embryos using fluorescein differential display: comparison of culture environments. Biology of reproduction 2001; 64: 30–35.

32. Ho, Y., Wigglesworth, K., Eppig, J. J., Schultz, R. M. Preimplantation development of mouse embryos in KSOM: augmentation by amino acids and analysis of gene expression. Molecular reproduction and development 1995; 41: 232–238.

33. Tsukiyama, T., Asano, R., Kawaguchi, T., Kim, N., Yamada, M., Minami, N., Ohinata, Y., Imai, H. Simple and efficient method for generation of induced pluripotent stem cells using piggyBac transposition of doxycycline-inducible factors and an EOS reporter system. Genes to Cells 2011; 16: 815–825.

34. Fan, X., Petitt, M., Gamboa, M., Huang, M., Dhal, S., Druzin, M. L., Wu, J. C., Chen-Tsai, Y., Nayak, N. R. Transient, inducible, placenta-specific gene expression in mice. Endocrinology 2012; 153: 5637–5644.

35. Shikata, D., Yamamoto, T., Honda, S., Ikeda, S., Minami, N. H4K20 monomethylation inhibition causes loss of genomic integrity in mouse preimplantation embryos. Journal of Reproduction and Development 2020; 66: 411–419.

36. Livak, K. J., Schmittgen, T. D. Analysis of relative gene expression data using real-time quantitative PCR and the 2(-Delta Delta C(T)) Method. *Methods (San Diego*, Calif.) 2001; 25: 402–408.

37. Xiang, J., Chao, D. T., Korsmeyer, S. J. BAX-induced cell death may not require interleukin 1β-converting enzyme-like proteases. Proceedings of the National Academy of Sciences of the United States of America 1996; 93: 14559–14563.

38. Riedl, S. J., Shi, Y. Molecular mechanisms of caspase regulation during apoptosis. Nature Reviews Molecular Cell Biology 2004 5:11 2004; 5: 897–907.

39. Slee, E. A., Harte, M. T., Kluck, R. M., Wolf, B. B., Casiano, C. A., Newmeyer, D. D., Wang, H. G., Reed, J. C., Nicholson, D. W., Alnemri, E. S., Green, D. R., Martin, S. J. Ordering the cytochrome c-initiated caspase cascade: Hierarchical activation of caspases-2,-3,-6,-7,-8, and -10 in a caspase-9-dependent manner. Journal of Cell Biology 1999; 144: 281–292.

40. Benabdellah, K., Cobo, M., Muñoz, P., Toscano, M. G., Martin, F. Development of an All-in-One Lentiviral Vector System Based on the Original TetR for the Easy Generation of Tet-ON Cell Lines. PLoS ONE 2011; 6: e23734.

41. Michalec-wawiórka, B., Czapiński, J., Filipek, K., Rulak, P., Czerwonka, A., Tchórzewski, M., Rivero-müller, A. An improved vector system for homogeneous and stable gene regulation. International Journal of Molecular Sciences 2021; 22: 5206.

